# A modiolar-pillar gradient in auditory-nerve dendritic length: a novel post-synaptic contribution to dynamic range?

**DOI:** 10.1101/2024.11.04.621861

**Authors:** Serhii Kostrikov, Jens Hjortkjaer, Torsten Dau, Gabriel Corfas, Leslie D. Liberman, M. Charles Liberman

## Abstract

Auditory-nerve fibers (ANFs) from a given cochlear region can vary in threshold sensitivity by up to 60 dB, corresponding to a 1000-fold difference in stimulus level, although each fiber innervates a single inner hair cell (IHC) via a single synapse. ANFs with high-thresholds also have low spontaneous rates (SRs) and synapse on the side of the IHC closer to the modiolus, whereas the low-threshold, high-SR fibers synapse on the side closer to the pillar cells. Prior biophysical work has identified modiolar-pillar differences in both pre- and post-synaptic properties, but a comprehensive explanation for the wide range of sensitivities remains elusive. Here, in guinea pigs, we used immunostaining for several neuronal markers, including Caspr, a key protein in nodes of Ranvier, to reveal a novel modiolar-pillar gradient in the location of the first ANF heminodes, presumed to be the site of the spike generator, just outside the sensory epithelium. Along the cochlea, from apex to base, the unmyelinated terminal dendrites of modiolar ANFs were 2 - 4 times longer than those of pillar ANFs. This modiolar-pillar gradient in dendritic length, coupled with the 2 - 4 fold smaller caliber of modiolar dendrites seen in prior single-fiber labeling studies, suggests there could be a large difference in the number of length constants between the synapse and the spike initiation zone for low- vs high-SR fibers. The resultant differences in attenuation of post-synaptic potentials propagating along these unmyelinated dendrites could be a key contributor to the observed range of threshold sensitivities among ANFs.

## Introduction

Auditory nerve fibers (ANFs), comprising the primary sensory neurons of the auditory pathway, are bipolar, with a peripheral axon projecting towards the hair cells in the organ of Corti, and a central axon projecting to the cochlear nucleus in the brainstem (Spoendlin, 1969; Spoendlin, 1972). The great majority of these ANFs, the co-called type-I neurons, are myelinated throughout much of their course, including most of the peripheral axon traversing the osseous spiral lamina (Figure 1). Each peripheral axon loses its myelin near the habenula, where it enters the organ and Corti and typically forms a synaptic connection with a single inner hair cell (IHC) via an unmyelinated dendritic process (Liberman, 1980). At this synaptic contact, there is an active-zone complex with a patch of AMPA-type glutamate receptors in the ANF membrane (Matsubara et al., 1996) positioned opposite a small pre-synaptic ribbon in the IHC with associated voltage-gated Ca^++^ channels in the IHC membrane (Frank et al., 2010).

**Figure 1:**
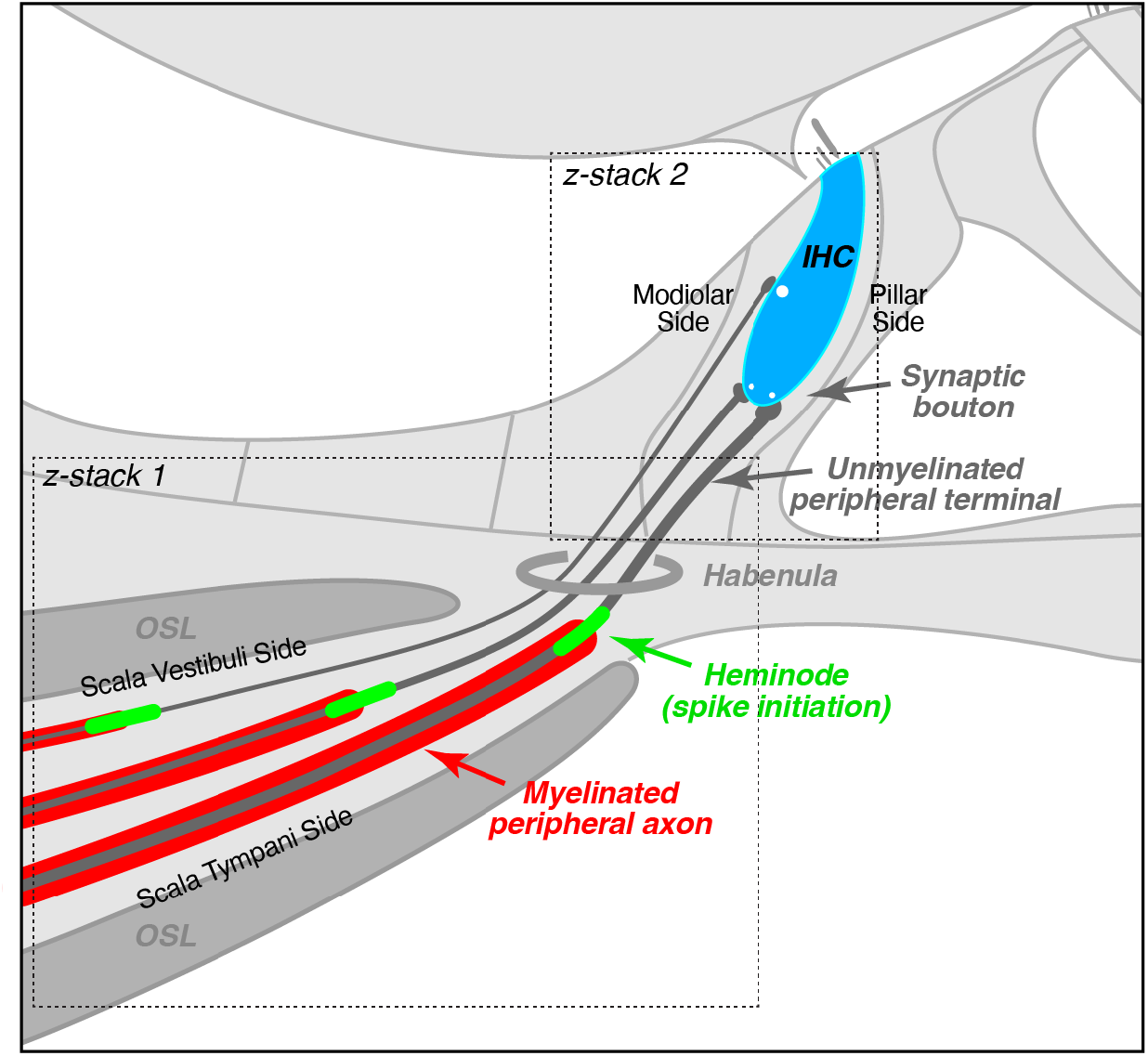
Schematic illustrating the peripheral projections of auditory nerve fibers from the ends of the myelinated peripheral axons in the osseous spiral lamina to the synaptic terminal in contact with the inner hair cell. The dashed squares illustrate how the composite region of interest was tiled via acquisition of two partially overlapping z-stacks.

Reflecting their punctate peripheral innervation, ANFs are exceedingly sharply tuned, faithfully mirroring the high degree of mechanical frequency selectivity at each position along the spiraling cochlear sensory epithelium (Kiang et al., 1965). However, among the fibers from a given cochlear location, i.e. tuned to the same frequency, there can be a range of threshold sensitivity of up to 60 dB, corresponding to a 1000 fold difference in stimulus sound pressure (Liberman, 1978). Serial-section ultrastructure of cat cochleas in the 1980’s showed that ANF dendrites synapsing on the modiolar side of the IHCs (Figure 1) were thinner and contained many fewer mitochondria compared with those on the pillar side (Liberman, 1980). This anatomical organization was hypothesized to reflect the spatial segregation of high-threshold fibers, which also showed very low rates of spontaneous discharge, from low-threshold fibers with much higher spontaneous rates (SRs). Subsequent intracellular labeling experiments in cat (Liberman, 1982) and guinea pig (Tsuji and Liberman, 1997) confirmed this hypothesis. However, the difference in mitochondrial content is likely an effect of the SR differences, rather than a cause. Subsequent *in vitro* studies in mice using calcium imaging of IHCs and/or patch-clamp recordings from the unmyelinated terminals of ANFs have noted a variety of pre- and post-synaptic differences between the modiolar and pillar sides of the IHCs. These synaptic differences could contribute to the range of sensitivities among the ANFs that contact them (for review, see (Moser et al., 2023)), however, none seems of a magnitude sufficient to explain the full 60 dB dynamic range.

Here, we used confocal imaging and immunostaining in the guinea pig cochlea, combined with morphometric analysis, to map the locations of the putative ANF spike initiation zones, i.e. the first heminodes in the osseous spiral lamina (Kim and Rutherford, 2016), relative to their IHC synapses. We found that the unmyelinated terminals of ANFs on the modiolar sides of the IHCs are up to 4 times longer than those on the pillar side. This modiolar-pillar gradient in dendritic length, coupled with the 2-fold smaller mean diameter of modiolar terminals (Liberman and Oliver, 1984; Tsuji and Liberman, 1997), could produce large differences in the attenuation of EPSPs as they propagate from the IHC synapse to the spike generator and thus may be an important contributor to the observed differences in threshold sensitivity and SR.

## Methods

### A. Tissue fixation, immunostaining and image acquisition

Animals were female albino guinea pigs of the Hartley strain, obtained at ∼250 g. At ages of 30 – 50 days, animals were deeply anesthetized with urethane (900 mg/kg) followed by fentanyl and haloperidol (0.15 and 10 mg/kg, respectively), then tissue was fixed by intravascular perfusion of 4% paraformaldehyde following a saline wash-out. Cochleae were flushed through the scalae with the same fix and then post-fixed for 2 hrs, decalcified for 2–3 weeks in 0.12 M EDTA, and dissected into roughly 11 pieces, each containing a fraction of the spiraling sensory epithelium with the osseous spiral lamina attached.

Cochlear pieces were permeabilized by freezing on dry ice in 30% sucrose and blocked for 1 hr at 22°C in PBS with 1% Triton X and 5% normal horse serum (NHS). Tissue was then incubated for 3 nights at room temperature on a shaker in the following primary antibodies, diluted in PBS with NHS and Triton X (both at 1%): 1) mouse isotype IgG2a anti-Tubulin beta 3 (TUJ1, 1:1000, BioLegend #801201), 2) chicken anti-neurofilament H (NFH, 1:500, Millipore #AB5539), 3) mouse isotype IgG1 anti-C-terminal binding protein 2 (CtBP2, 1:100, BD Transduction Laboratories #612044), 4) mouse isotype IgG1 anti-Caspr/paranodin/neurexin IV (Caspr, 1:10, NeuroMab clone #K65/35, deposited to the Developmental Studies Hybridoma Bank by J.S. Trimmer), 5) rabbit anti-myosin VIIa (Myo7a, 1:100, Proteus BioSciences #25–6790), and goat anti-choline acetyltransferase (ChAT, 1:200, Millipore #AB144P). After washing in PBS, the pieces were incubated first in biotinylated Fab Fragment donkey anti-goat diluted in 1% NHS and Triton X in PBS for 1 hr at 37 deg C (1:200, Jackson ImmunoResearch #705-067-003), and then twice for 1 hr each at 37°C in the following secondaries, diluted in same: (1) Alexa Fluor 488 Fab Fragment goat anti-mouse(IgG2a) (1:500, Jackson ImmunoResearch #115-547-186), (2) Alexa Fluor 488 goat anti-chicken (1:1000, ThermoFisher #A11039), (3) Alexa Fluor 647 goat anti-mouse(IgG1) (1:200, ThermoFisher #A21240), (4) Pacific Blue goat anti-rabbit (1:200, ThermoFisher #P10994), and (5) Pacific Blue, streptavidin conjugate (1:200, ThermoFisher #S11222). Finally, the tissue was incubated for 30 min at room temperature on a shaker in CellMask Orange Plasma Membrane Stain (1:1000, ThermoFisher #C10045), diluted in PBS with 0.3% Triton X. The pieces were rinsed in PBS after each step and mounted in VectaShield with basilar membrane up and hair cells down.

Low-power images of the myosin channel in each microdissected piece were obtained with a 4x objective on a Nikon E800 epifluorescence microscope. These images were then arranged sequentially in a montage to measure the cumulative distance along the cochlear spiral using a custom ImageJ plug-in that superimposes frequency correlates onto the imaged spiral by application of the cochlear map specific to the guinea pig (Tsuji and Liberman, 1997).

The organ of Corti was imaged using a Leica confocal microscope (SP8) equipped with a 63x glycerol immersion objective (N.A. 1.3) at a digital zoom of 2.4, applying the Leica deconvolution algorithm to enhance resolution. The resulting z-stacks had pixel sizes of 48 nm in the x and y dimensions and 300 nm in the z dimension. In each ear (n=3), at each of 8 octave-spaced cochlear frequencies (0.25 to 32.0 kHz), two contiguous fields (comprising 8-9 IHCs per field) were imaged. For each field, the region of interest was tiled by first obtaining a z stack through the peripheral axons in the osseous spiral lamina, habenula and inner spiral bundle, followed by a second, partially overlapping, z-stack through the inner spiral bundle and inner hair cell area of exactly the same longitudinal position along the cochlear spiral (See Figure 1). Tiling of the two separate stacks was necessary because the fluorophores of interest differed between the peripheral axon region and the inner hair cell region.

### B. Histological and Morphometric analysis

The numbers and volumes of pre-synaptic ribbons were analyzed with Amira^®^ software (version 2019.4, Thermoscientific) using the *connected components* feature to automate the identification of CtBP2-positive puncta in three-dimensional space, and the notation of their locations, sizes and numbers. After their automatic segmentation, the clouds of ribbon points were visually cross-checked against maximum projections of the native z-stacks to identify and correct false negatives and false positives.

To locate the heminodes, the images from the Caspr channel underwent transformation from 16 to 8 bits. They were then upscaled by a factor of 2 in each dimension using bilinear interpolation and subjected to 3D Gaussian filtering (x, y, z σ = 2) to smooth the image without substantial loss of resolution. The resulting images were deblurred by subtracting a copy of the image, which underwent 3D Gaussian filtering (σ of x,y,z = 5). Then the images were binarized by thresholding (pixel intensity: 30–255) and the binary mask underwent maximum filtering (kernel size = 1) in the z plane.

The images from the myelin (Cellmask) channel were also upscaled to match the dimensions of their Caspr counterparts. For obtaining raw myelination masks, the images underwent Gaussian filtering (σ of x,y,z = 5) and subsequent thresholding (pixel intensity: 3000–65535). A binary mask of the myelin border was obtained by subtracting myelination mask from its copy, which underwent maximum filtering (kernel = 7) in the y axis (parallel to fiber direction). The resulting mask was inverted and subtracted from the mask of the nodes acquired from the Caspr channel to obtain initial indications of the myelination endpoints. These masks underwent further processing: 3D maximum filtering (kernel size in x, y, z = 1) and Gaussian filtering (σ values: x=12, y=10, z=3) with subsequent thresholding (pixel intensity: 18–255). All the above-mentioned operations were performed in Fiji (version 1.53t).

Processed myelination endpoint masks were then visually examined, and all the detection inaccuracies were corrected. Efferent fibers (indicated by the ChAT staining) were excluded from the analysis, as were a small number of true nodes, i.e. locations at the junction of the first and second myelinated segments. Final endpoint masks underwent label analysis in Amira^®^ 3D (version 2021.1, Thermoscientific) to extract the coordinates of the centers of gravity of the discrete objects.

The manually curated set of y,z coordinates of the ribbons and heminodes in each pair of z-stacks were then transformed into new coordinate systems based on the modiolar-pillar polarity of the IHCs or the vestibuli-tympani polarity of the osseous spiral lamina, respectively, using a custom LabVIEW program (Hickman et al., 2020). This program required user input to define a pair of lines, intersecting in the middle of the habenula, as judged from the maximum zy projection of each z-stack: one of these lines bisected the subnuclear portion of the IHCs into pillar vs. modiolar halves, and the other bisected the osseous spiral lamina into its scala tympani vs. scala vestibuli facing sides. These bisectors became the transformed y axes for ribbons and heminodes, respectively, and a pair of x axes was created perpendicular to each running through the midpoint of the habenula. See text for further details.

### D. Statistical analysis

Regression analyses and t-tests were carried out in Excel.

## Results

To study the unmyelinated terminal dendrites of ANFs, we microdissected and immunostained the cochleas of young normal guinea pigs. Confocal z-stacks were acquired at each of eight cochlear frequency regions, spanning the entire 21 mm spiraling epithelium in octave steps from the 0.25 kHz region (∼90% of the distance from the base) to the 32.0 kHz region (∼10% of the distance from the base). As illustrated in Figure 1, the z-stacks were positioned to capture the peripheral ends of type-I ANFs, extending from their synapses with the IHC to the first heminode, where the spike initiation zone is presumed to be located (Kim and Rutherford, 2016) and where the fibers become myelinated (Liberman, 1980). We used a lipophilic fluorophore (Cellmask®) to label the myelin sheaths and antibodies to Contactin Associated Protein 1 (Caspr) to label the paranodal and juxtaparanodal regions of the nodes of Ranvier (Hossain et al., 2005; Peles et al., 1997). Additionally, we used antibodies to C-terminal Binding Protein 2 (CtBP2) to label synaptic ribbons (Khimich et al., 2005) and Choline Acetyl Transferase (ChAT) to label olivocochlear efferent fibers (Eybalin and Pujol, 1987) allowing us to exclude them from the analysis, since their peripheral axons can also be myelinated in the osseous spiral lamina.

As shown in the representative maximum projections in Figure 2, the Caspr-positive heminodes (green) appear at the distal tips of the myelinated fibers (red). Manual examination by scrolling single z-slice views through this (and every other) stack verified that every myelin sheath terminated with a heminode, and every heminode was located at the end of the myelin sheath. Occasionally, thicker Caspr-positive profiles were observed in a characteristic doublet formation: single-slice scrolling confirmed that these doublet profiles correspond to first nodes, since the myelin sheath continued on both ends. Prior studies in mouse confirm that first nodes typically appear >50 μm from the habenula (Hossain et al., 2005; Kim and Rutherford, 2016; Wan and Corfas, 2017), and our z-stacks span only about the distalmost 40 μm of the ANFs (Figure 2).

**Figure 2:**
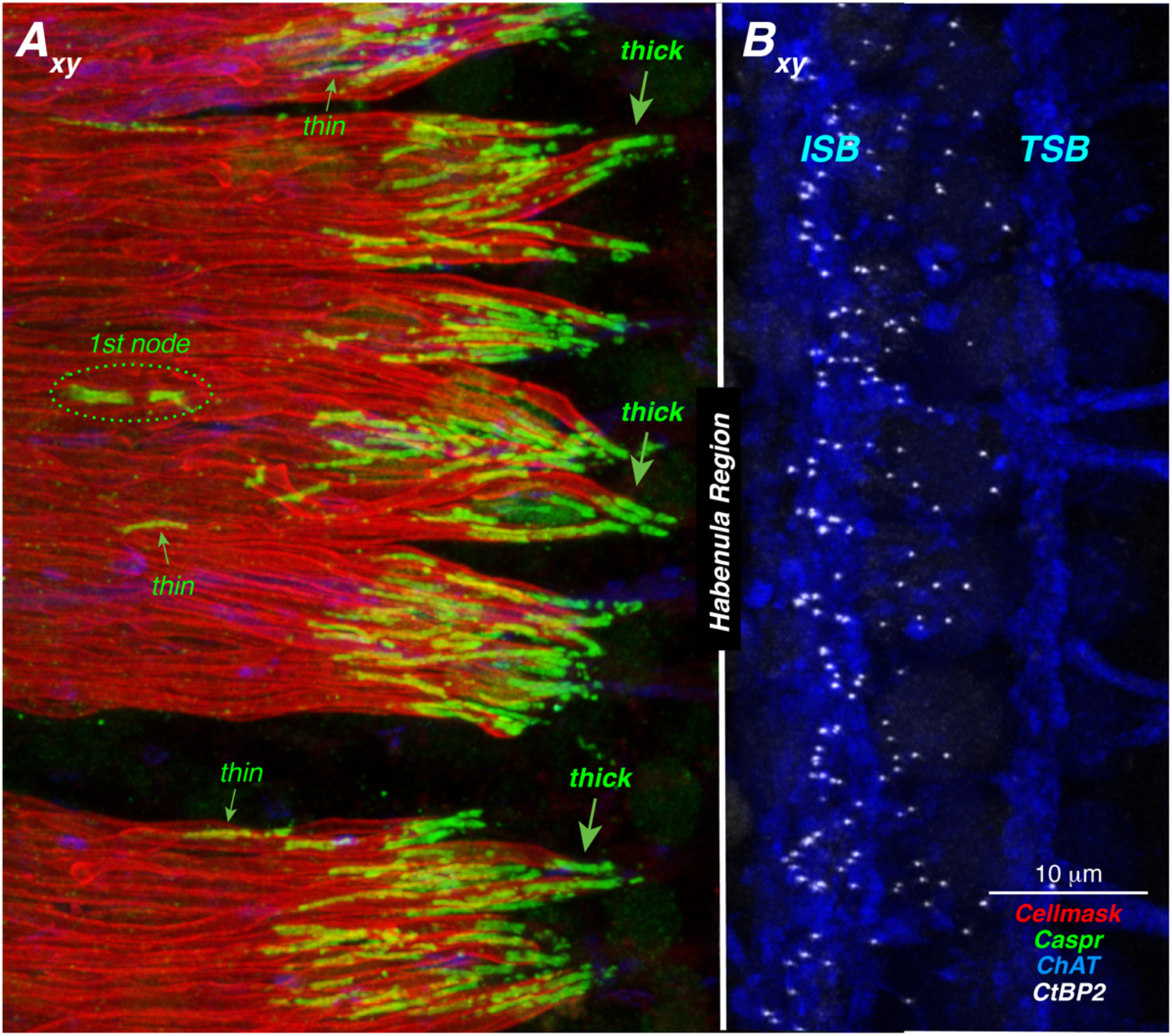
Representative confocal images showing the immunostaining used to reveal (**A**_**xy**_) the myelin sheaths (Cellmask; red) and initial heminodes (Caspr; green) of auditory nerve fibers in the osseous spiral lamina, as well as (**B**_**xy**_) the presynaptic ribbons (CtBP2; white) in the inner hair cells and the olivocochlear efferent innervation (ChAT; blue). Each image is a maximum projection onto the z plane of a confocal z-stack from the 8.0 kHz region. The two images are in correct registration as they would appear when viewed in the acquisition (xy) plane, although they were acquired as separate z-stacks, digitally shifting the region of interest in the radial direction, parallel to the nerve-fiber trajectories, as schematized in Figure 1.

The images in Figure 2 also reveal that heminodes are present at variable distances from the habenula region, ranging from less than 5 μm to more than 40 μm. Visual examination of the profiles also suggests that the heminodal profiles located farther from the habenula are also thinner in caliber. Similarly, the synaptic ribbons (i.e. the CtBP2-positive puncta in Figure 2B_xy_) are located at varying distances from the habenula, ranging from less than 5 μm to greater than 20 μm. Viewing these same z-stacks in the zy plane (Fig. 3) provides a more accurate perspective of the distances involved. It also reveals complementary gradients, whereby the heminodes farthest from the habenula tend to be located on the scala vestibuli side of the osseous spiral lamina, while the synapses farthest from the habenula tend to be on the modiolar side of the IHC. Prior studies have suggested that dendrites synapsing on the modiolar side connect with peripheral axons on the scala vestibuli side of the lamina (Kawase and Liberman, 1992), indicating that high-threshold, low-SR fibers generally have longer unmyelinated dendrites than the low-threshold, high-SR fibers.

**Figure 3:**
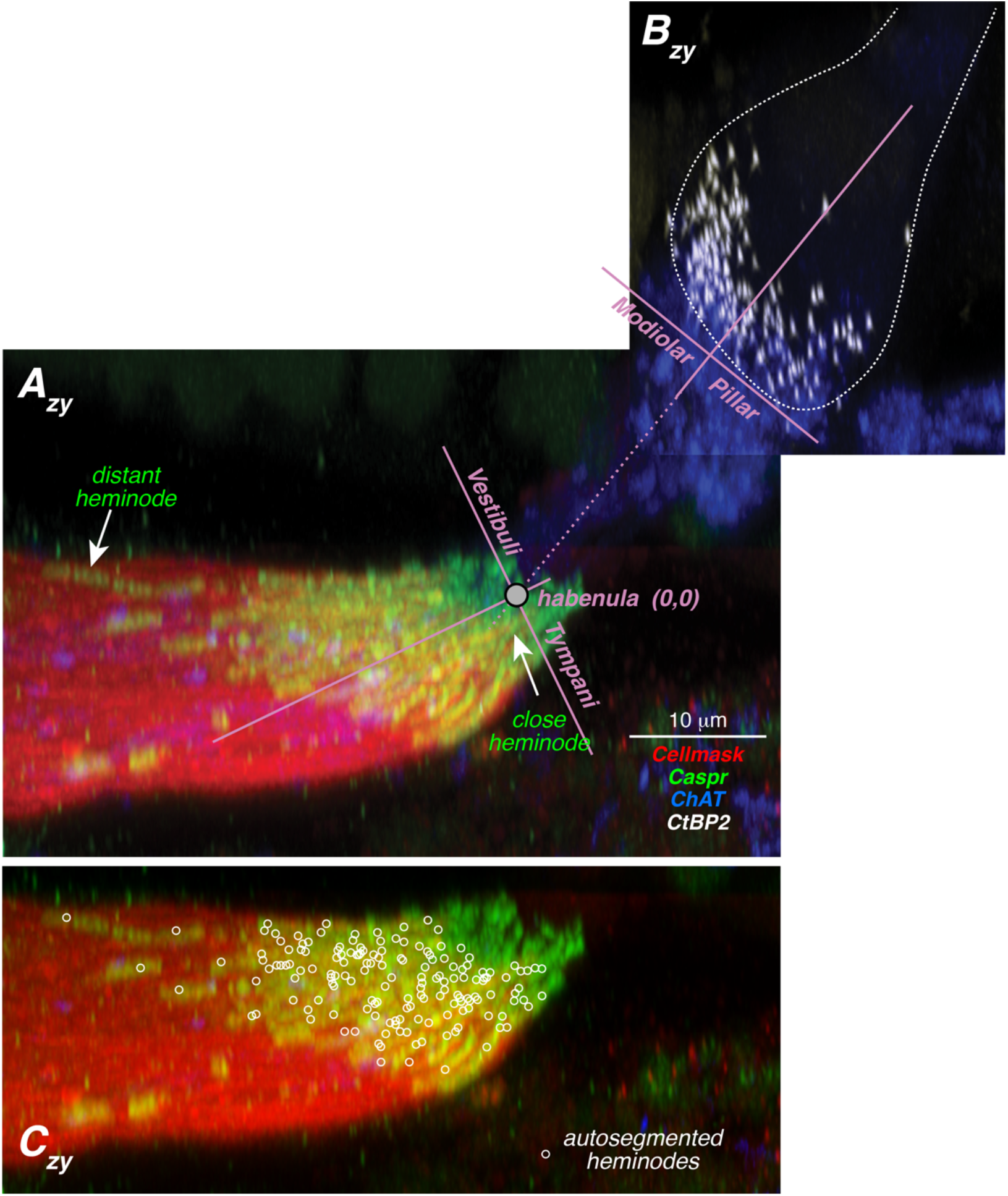
Axis transformations used to quantify the locations of ANF heminodes (**A**_**zy**_, **C**_**zy**_) and IHC presynaptic ribbons (**B**_**zy**_) in the present study. Each image is the maximum projection onto the x plane of the same confocal z-stacks shown in Figure 2. Superimposed on the tiled images in **A**_**zy**_ and **B**_**zy**_ are the two sets of axes used to express the relative locations of heminodes and synaptic ribbons along the vestibuli-tympani or modiolar-pillar axis, respectively. The two sets of axes share the same origin and are placed in the middle of the habenula (0,0). The locations of the heminodes in this case are shown by the open circles in **C**_**zy**_.

To quantitatively assess these gradients, we developed workflows to autosegment the ribbons and heminodes. Techniques for segmentation and 3D mapping of synaptic ribbons were already well established in prior studies (Hickman et al., 2020; Liberman et al., 2015) and their accuracy is easily verify by rotating and comparing surface renderings of the autosegmented ribbon puncta with maximum projections of the confocal images. Our heminodal autosegmentation workflow was designed to identify the distal tip of each myelin sheath, specifically the point of overlap between each myelin mask and its associated heminode mask. The results for one z-stack are shown in Figure 3C_zy_, where heminode positions are indicated by white circles. In regions where the heminodes are scarce, the accuracy of the workflow is easy to assess, and this type of superposition was performed in all ears, for both zy and xy projections. As a further check on accuracy, a different observer took virtual cross-sections through the osseous spiral lamina, as illustrated in Figure 4A_zy_, and manually counted the ANF peripheral axons in slices such as the one shown in Figure 4B_xz_. The agreement between heminodal counts and fiber counts was very close (Fig. 4D). Note the caliber gradient in these virtual cross sections, with thinner fibers on the scala vestibuli side of the lamina (Fig. 4B_xz_). When slicing closer to the habenula (Fig. 4C_xz_), many of these thinner fibers have lost their myelin, leaving only their heminodes visible. Immunostaining for a cholinergic marker (ChAT) allowed us to exclude olivocochlear efferent axons from the present analysis (blue arrowhead in Figure 4B_xz_).

**Figure 4:**
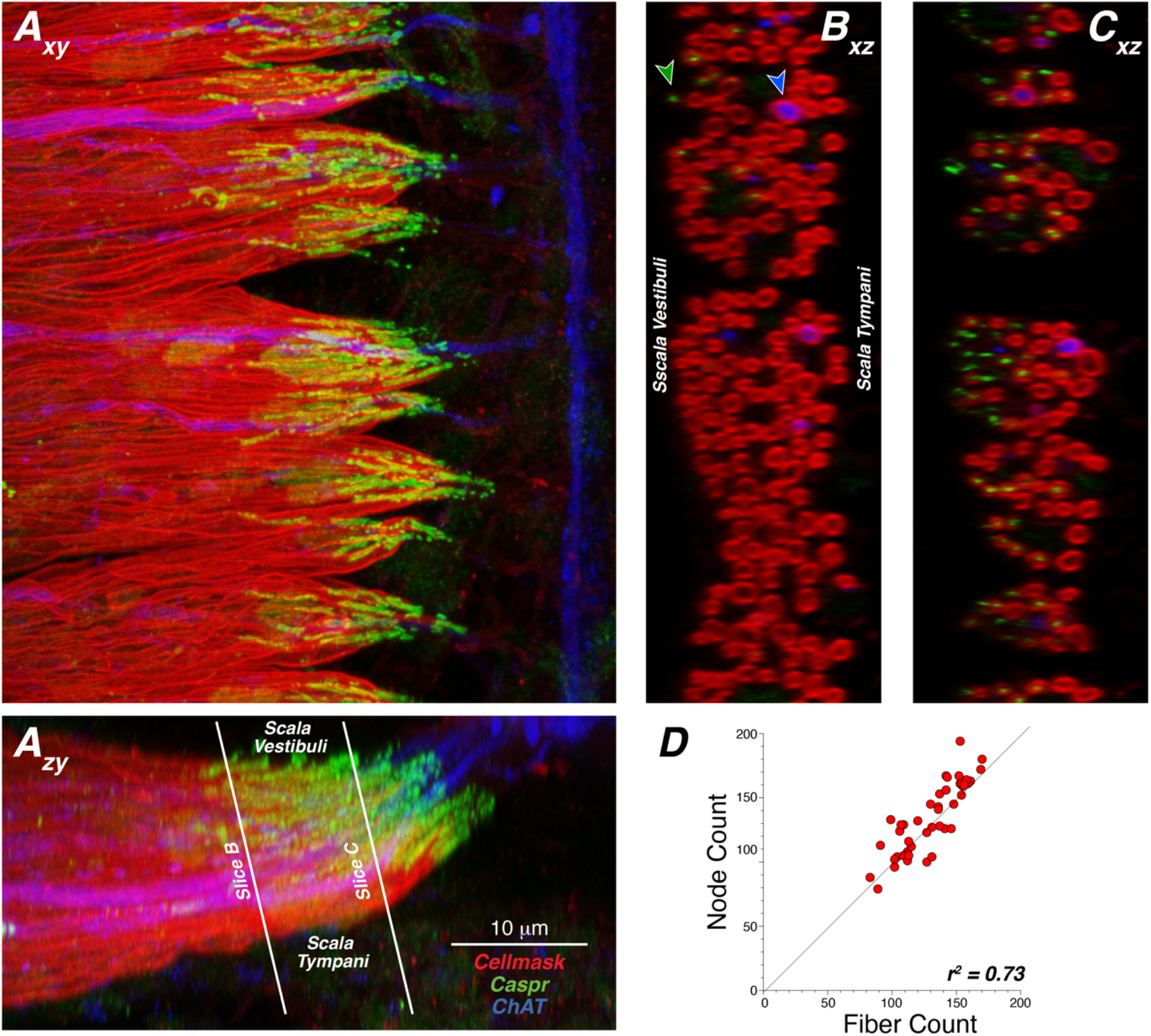
Automated counts of heminodes are closely correlated with manual counts of myelinated fibers in the same z-stacks. **A**_**xy**_ and **A**_**zy**_ are maximum projections of a confocal z-stack from the 5.6 kHz region, onto the z and x planes, respectively. **B**_**xz**_ and **C**_**xz**_ are virtual slices through this z-stack taken at the locations, and with the orientations, shown in **A**_**zy**_. A section through a heminode captured distal to the loss of myelination is shown at the green-fill arrowhead. A myelinated olivocochlear efferent is seen opposite the blue-fill arrowhead. The scatterplot in **D** compares the manual fiber counts taken from images such as **B**_**xz**_ with the automated heminode counts such as those shown in **Figure 3C**_**zy**_. Each point shows data from a different z-stack. The dashed diagonal is the reference line for a unity ratio.

To further quantify the complementary spatial gradients of heminodes and synaptic ribbons, and to estimate the lengths of ANF peripheral dendrites, we defined two new sets of axes for each z-stack. As shown in Figure 3A_zy_, each axis had its origin (0,0) at the center of the habenula, the area through which all fibers must pass. One pair of axes was visually adjusted to bisect the osseous spiral lamina, particularly in the region close to the habenula, while the other pair was set to bisect the synaptic ribbon cloud at the basolateral pole of the IHC.

This axis transformation allowed positional data to be superimposed across different z-stacks. As previously described (Hickman et al., 2020), synaptic ribbons cluster at the basolateral pole of the IHC (Fig. 5A,B), and the ribbons on the modiolar side tend to be larger (Fig. 5C). In the present study this size gradient was consistently present, and highly significant, across all cochlear regions evaluated (Fig. 5D). There is also a ribbon size gradient along the habenular cuticular axis, expressed here as distance from the basal end of the IHC (Fig. 5E). That gradient was highly significant, according to simple linear regression, at all cochlear regions except the basal and apical extremes (Fig. 5F).

**Figure 5:**
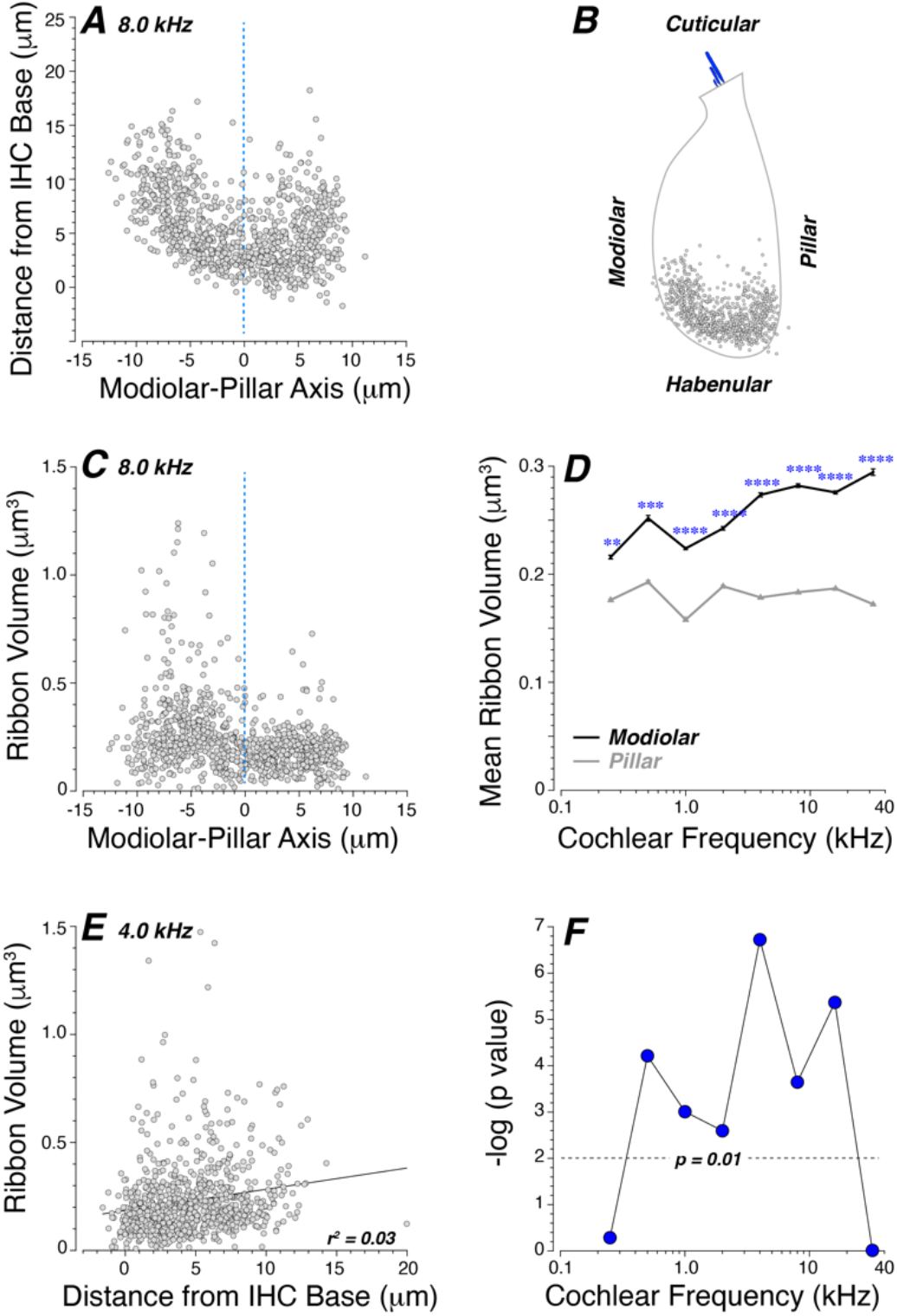
Synaptic ribbons show size gradients along both the modiolar-pillar and the habenular-cuticular axes. Panel **A** shows superimposed locations of pre-synaptic ribbons, as extracted from 2 z-stacks from each of 3 ears at the 8 kHz location (a total of 897 ribbons and 49 IHCs), and after transformation to a hair-cell centric set of axes, as illustrated in Figure 3. Panel **B** shows the orientation of the ribbon cloud with respect to an approximate outline of the IHC silhouette. Panel **C** shows the volumes of these 8 kHz ribbons as a function of their modiolar-pillar position. Panel **D** shows the mean modiolar vs. pillar volumes at each of 8 cochlear locations (± SEMs) based on data from 2 z-stacks from each of 3 ears, with significance values shown by asterisks (* p < 0.05; ** p < 0.01; *** p < 0.001; **** p << 0.0001. Panel **E** shows the volumes of presynaptic ribbons as a function of their position along the habenular-cuticular axis, expressed as distance from the IHC base after the same axis transformation shown in Figure 3. Data are from 2 z-stacks from each of 3 ears at the 4.0 kHz region (a total of 823 synapses and 46 IHCs). The regression line and its r^2^ value are also shown. Panel **F** shows the significance values for the regressions at all 8 cochlear frequency regions between ribbon volume and distance from IHC base. Significance values are expressed as -log(p).

The spatial gradients of heminodes and synapses are summarized in Figure 6 for all eight cochlear regions in each of the three cases analyzed. With respect to heminodal position, the data show that ANFs on the scala vestibuli side of the osseous spiral lamina had longer peripheral dendrites, i.e. became myelinated farther from the habenula, compared to ANFs on the scala tympani side. These length differences ranged from at least 20 μm to at most 40 μm. Regression analysis revealed that this length gradient was highly significant (p< 0.001) for all data shown in the scatterplots in Figure 6 except for the 0.25 kHz case from the top row. At 0.5 kHz and above, the p values for the regression analyses were all << 0.0001. The inferred gradient of dendritic lengths within the osseous spiral lamina is compounded with a complementary spatial pattern within the organ of Corti, wherein the synaptic ribbons with the largest distances from the habenula tend to be located at the most modiolar extremes. Thus, as illustrated for one case at the 8 kHz region, dendritic lengths can vary systematically with synaptic position by a factor of 4, from roughly 80 μm at the modiolar extreme to only 20 μm on the pillar side. It is notable that these ranges for dendritic distances vary little across the entire cochlear spiral.

**Figure 6:**
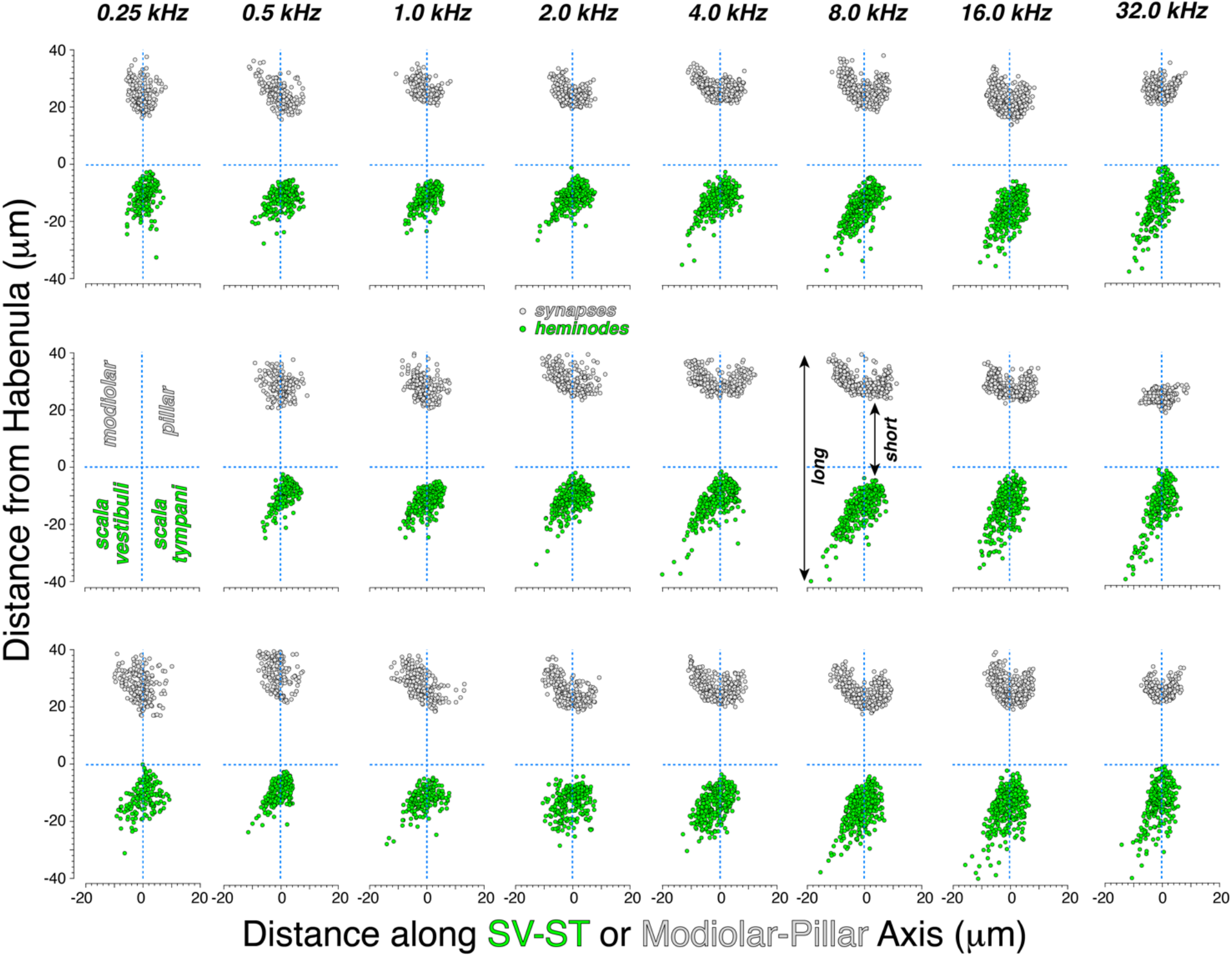
The unmyelinated terminals of ANFs innervating the modiolar side of the IHC can be up to 4 times as long as those on the pillar side. Each of the 23 scatterplots summarizes the distances of presynaptic ribbons (black) and ANF heminodes (green) from the habenula, as a function of their position along the modiolar-pillar axis of the IHCs or the vestibuli-tympani sides of the OSL (see Figure 3). Cochlear locations are as indicated by the column labels; each row show data from a different ear. Distance from the habenula is computed from each transformed (x,y) coordinate to the respective axis origin (0,0).

## Discussion

### A. Mechanisms underlying sensitivity differences among ANFs

The dynamic range of the auditory system is remarkable. The difference in sound pressure between threshold levels and traumatic levels is ∼120 dB, corresponding to a million-fold range in amplitude of the sound waves (Geisler, 1998), yet humans can detect loudness differences of less than 1 dB across most of this dynamic range (Viemeister and Bacon, 1988). Some of this dynamic range is likely mediated by the spread of excitation along the mechanically tuned cochlear epithelium as sound pressures increase (Kiang and Moxon, 1974); however, different ANFs innervating the same cochlear region can vary in threshold sensitivity by > 60 dB (Liberman, 1978).

Thresholds in ANFs are highly correlated with spontaneous discharge rate (SR), i.e. the background spike rate in the absence of acoustic stimulation (Liberman, 1978). SR can vary significantly in different fibers, from near 0 to over 100 sp/sec (Kiang et al., 1965). In cat, where this correlation has been most extensively studied, the relation between threshold and SR resembles a step function rather than a continuum: “high-SR” fibers (SR > 20 sp/sec) have similar low thresholds and are ∼10 dB more sensitive than medium-SR fibers (0.5 < SR < 20 sp/sec), which, in turn, are 10 – 50 dB more sensitive the low-SR group (0 < SR < 0.5 sp/sec) (Liberman, 1978). The relationship between SR and threshold is fundamentally similar in guinea pigs (Tsuji and Liberman, 1997) and mice (Taberner and Liberman, 2005).

Serial-section ultrastructure studies have shown that the unmyelinated peripheral terminals of ANFs are almost always unbranched. Each fiber contacts a single IHC, via a single terminal bouton containing a discrete active zone of thickened post-synaptic membrane, opposite 1 - 2 pre-synaptic ribbons surrounded by a halo of synaptic vesicles (Liberman, 1980; Spoendlin, 1969). Modiolar fibers are thinner and contain fewer mitochondria than pillar-side fibers, suggesting the former might have lower SR and higher thresholds (Liberman, 1980). Intracellular labeling experiments in cat (Liberman, 1982) and then guinea pig (Tsuji and Liberman, 1997) confirmed the functional significance of this spatial segregation. As predicted, low- and medium-SR fibers synapsed exclusively on the side of the IHC facing the modiolus, while most high-SR fibers synapsed on the side of the IHCs facing the pillar cells (Fig 1). These labeling experiments also demonstrated that low-SR fibers are thinner than high-SR fibers, in their unmyelinated peripheral terminals (Liberman, 1982; Tsuji and Liberman, 1997) as well as in their myelinated peripheral and central axons (Liberman and Oliver, 1984).

The legacy physiological data suggesting that ANFs comprise three discrete groups have been corroborated in mice by cluster analysis of gene expression in ANF cell bodies (Shrestha et al., 2018; Sun et al., 2018). Furthermore, the spatial segregation of terminals on the IHC has been confirmed using immunohistochemistry and/or *in situ* hybridization for cluster-specific markers (Shrestha et al., 2018); and *in vitro* recording from the terminal boutons of genetically labeled fibers has verified that the modiolar-side clusters correspond to fibers with low- and medium SRs (Siebald et al., 2023).

Although the polarization of the IHC with respect to ANF SR groups is now definitively established using *in vivo* and *in vitro* approaches in several mammalian species, the mechanism(s) underlying the differences in threshold sensitivity remain unclear. There is no reason to think that the transmembrane potential, whether at rest or during sound stimulation, differs between the two sides of the hair cell. Nevertheless, as reviewed elsewhere (Moser et al., 2023), there are systematic modiolar-pillar differences in the properties of the presynaptic active zones that shape the relation between transmembrane voltage and calcium influx as well as vesicle release (Ohn et al., 2016). Additionally, there are differences in the electrophysiological properties of the ANFs themselves that can affect excitability (Markowitz and Kalluri, 2020) and differences in the degree of olivocochlear innervation of the unmyelinated ANF dendrites (Liberman, 1980; Liberman et al., 1990) that can contribute to the creation of a sensitivity gradient.

### B. Dendritic length difference as a contributor to ANF sensitivity

The present study introduces a novel structural aspect of the systematic modiolar-pillar gradients in IHC innervation that could also contribute to the ANF sensitivity gradient. Here, we show that the unmyelinated ANF terminals innervating the modiolar side of the IHC can be up to 4 times longer than those on the pillar side. These distances are minimum estimates, since they are based on straight-line radial projections from the heminode to the center of the nearest habenular opening, and from the center of the habenula to the nearest synaptic ribbon. We know from single-fiber labeling of ANFs (Liberman, 1982; Liberman and Oliver, 1984; Tsuji and Liberman, 1997) and serial section ultrastructure (Hashimoto et al., 1990; Liberman, 1980; Spoendlin, 1972) that ANF peripheral projections take a fundamentally radial course from their synaptic termini to their heminodes in the osseous spiral lamina. However, intriguingly, an ultrastructural study showed that the thinnest modiolar dendrites often further increased their lengths by moving to a habenular opening offset from the one closest to the IHC (see Figure 1 in (Liberman, 1980)).

This systematic difference in dendritic lengths, combined with the known caliber difference between modiolar and pillar terminals (a 2-fold difference in mean diameter and a 4-fold diameter difference between the thickest and thinnest fibers (Tsuji and Liberman, 1997)), could have major effects on threshold sensitivity by creating a large difference in the degree to which EPSPs are attenuated between the synapse and the spike initiation zone.

Immunohistochemical studies of cochlear Na+ and K+ channels suggest that the spike initiation zone for ANFs is located at the first heminode, where the ANFs become myelinated (Kim and Rutherford, 2016). Thus, ANF terminals function as dendrites, and any post-synaptic potentials initiated at the synapse will be attenuated as they propagate to the spike initiation zone, due to the passive cable properties of the unmyelinated segment. The magnitude of this attenuation depends on the length constant, the distance over which EPSP amplitude decreases to ∼1/3 of its original value. Length constants decrease in proportion to the square root of fiber radius (Katz, 1966), which translates to a factor of 2 difference between the thinnest low-SR and the thickest high-SR fibers. Factoring in a 4-fold difference in terminal lengths for longest low-SR vs the shortest high-SR fibers (Figure 6), suggests there could be a difference of several length constants between the synapse and the spike initiation zone for a low- vs. a high-SR terminal.

Although length constants have not been measured for ANF terminals, the value for dendrites of principal cells in the medial superior olive is reported to be 76.4 μm, and these dendrites tend to be thicker than ANF dendrites, with diameters ranging from 1 – 3 μm (Winters et al., 2017) compared to 0.3 to 1.2 μm (Tsuji and Liberman, 1997). Thus, the length constant for ANF dendrites could well be shorter than the distances spanned between the synapse and the heminode. Of course, these ideas assume similar membrane properties for high- and low-SR dendrites, which may not be the case. For example, there could be differences in the type or number of voltage-activated K+ channels that could compensate for the differences in diameter and length. Relevant data on this do not exist for ANFs. However, it is tempting to speculate that the small number of exceptionally thin and distant heminodes, such as those highlighted in Figure 2, correspond to the relatively small population of low-SR fibers with exceptionally high (i.e. 60 dB) relative thresholds (Liberman, 1978). Although the few relevant studies of recordings from ANF unmyelinated terminals within the organ of Corti suggest that a high proportion of EPSPs (>80%) trigger action potentials (Rutherford et al., 2012; Siegel, 1992), it is possible that such recordings do not sample the very thin peripheral terminals of high-threshold, low-SR fibers.

Further evidence that dendritic length might be correlated with SR and threshold is seen in the gradients of synaptic ribbon sizes. We showed here that synaptic-ribbon volume increases with increasing synaptic distance from the base of the IHC (Figure 5E,F). Intracellular labeling experiments have shown that low-SR fibers are associated with larger synaptic ribbons than high-SR fibers (Merchan-Perez and Liberman, 1996), and numerous immunohistochemical studies have shown that modiolar-side ribbons are larger than pillar-side ribbons (Hickman et al., 2020; Liberman et al., 2011), as is also demonstrated here (Figure 5C,D).

It is also intriguing that dendritic lengths of ANFs don’t vary along the cochlear spiral (Figure 6), in contrast to virtually all other dimensional aspects of the organ of Corti, which grow monotonically from base to apex. For example, the width of the basilar membrane (Cabezudo, 1978) and the height of the OHC stereocilia (Yarin et al., 2014) both increase by a factor of 4. As shown in Figure 7, the IHC area does not scale with cochlear location in the same way that the OHC area does. This might be dictated by the need to maintain the same sensitivity gradient for ANF terminals across all best frequency regions, in contrast to the need to steadily vary mechanical tuning, which requires a steady increase in overall size, particularly for the electromotile OHCs.

**Figure 7:**
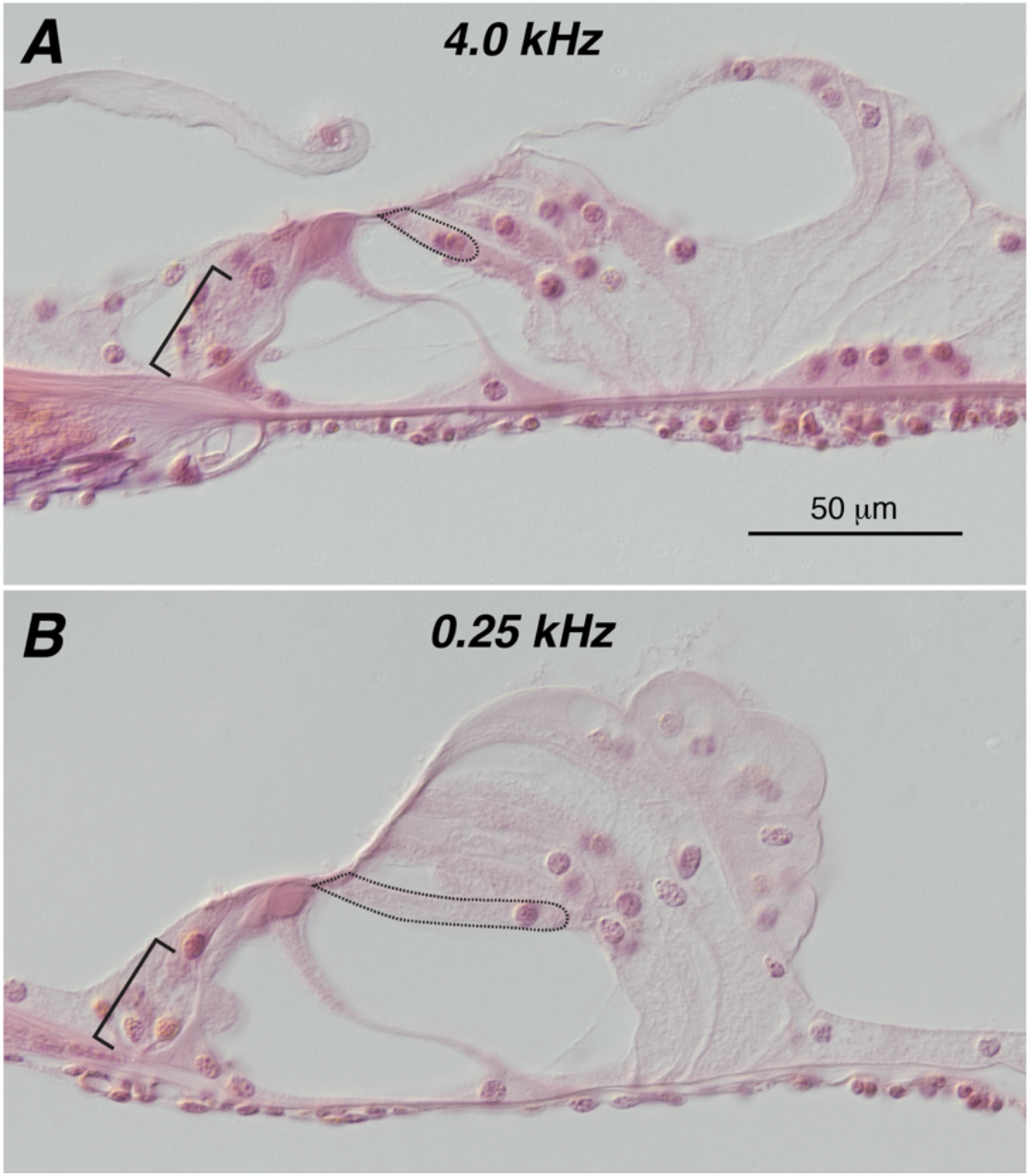
The dimensions in the IHC area are relatively invariant along the cochlear spiral, whereas the OHC area enlarges dramatically as best frequency decreases. The photomicrographs are from a normal adult guinea pig, processed by the celloidin technique and stained with hematoxylin and eosin (Merchant and Nadol, 2010). The brackets in the IHC area are the same length (30 μm) and, in both regions, span the distance from the basilar membrane to the bottom of the IHC nucleus. The OHCs in the first row are outlined in each image to show the apical-basal gradient in length, which would be even more dramatic if pictures were taken from basal extreme (45 kHz); however, this cochlear region (the cochlear “hook”) is cut in a very different plane making direct comparison more difficult.

## Acknowledgements

Research supported by a grant from the National Institute on Deafness and other Communicative Disorders (R01 DC 000188).

## Notes

**Conflict of Interest Statement:** The authors have no conflicts of interest or relevant financial relationships to disclose.

### Competing Interest Statement

The authors have declared no competing interest.

